# A New Highly Efficient Molecule for Both Optogenetic and Chemogenetic Control Driven by FRET Amplification of BioLuminescence

**DOI:** 10.1101/2023.06.26.545546

**Authors:** Andreas Björefeldt, Jeremy Murphy, Emmanuel L Crespo, Gerard G Lambert, Mansi Prakash, Ebenezer C Ikefuama, Nina Friedman, Tariq M Brown, Diane Lipscombe, Christopher I Moore, Ute Hochgeschwender, Nathan C Shaner

## Abstract

**Significance:** Bioluminescent optogenetics (BL-OG) offers a unique and powerful approach to manipulate neural activity both opto- and chemogenetically using a single actuator molecule (a LuMinOpsin, LMO).

**Aim:** To further enhance the utility of BL-OG by improving the efficacy of chemogenetic (bioluminescence-driven) LMO activation.

**Approach:** We developed novel luciferases optimized for Förster resonance energy transfer (FRET) when fused to the fluorescent protein mNeonGreen, generating bright bioluminescent (BL) emitters spectrally tuned to *Volvox* Channelrhodopsin 1 (VChR1).

**Results:** A new LMO generated from this approach (LMO7) showed significantly stronger BL-driven opsin activation compared to previous and other new variants. We extensively benchmarked LMO7 against LMO3 (current standard), and found significantly stronger neuronal activity modulation *ex vivo* and *in vivo*, and efficient modulation of behavior.

**Conclusions:** We report a robust new option for achieving multiple modes of control in a single actuator, and a promising engineering strategy for continued improvement of BL-OG.

## 1 Introduction

Understanding how specific cell types, circuits and brain areas are involved in generating perception, action and cognition requires approaches to experimentally manipulate spatiotemporal patterns of neural activity in genetically defined populations. At present, optogenetic and chemogenetic techniques constitute the major actuators used for this purpose, each with distinct strengths and limitations. Optogenetics notably offers the advantage of close to millisecond precision in the control of neural output [1, 2]. However, this approach requires invasive delivery of light, typically only practical to deliver in 1-2 locations in the brain or body. In comparison, chemogenetic approaches, including most notably designer receptors exclusively activated by designer drugs, DREADDs, provide sustained and minimally invasive modulation of neural populations distributed across much larger areas of the brain or body [3, 4]. To combine the strengths of above techniques and increase user flexibility, we have developed a bimodal strategy in which a luciferase-opsin fusion molecule (a luminopsin, LMO; Berglund et al., 2013; Berglund et al., 2016) integrates opto- and chemogenetic features in a single actuator [5]. With BL-OG, chemogenetic modulation of neural activity is achieved via delivery of a substrate, coelenterazine (CTZ), that is oxidized by the luciferase in the LMO, producing photons immediately adjacent to (within the same molecule) the optogenetic actuator. This approach allows the use of the same molecule for chemo- and optogenetic control at distinct times in the same animal, or individually [6–9]. As one example, Medendorp et al. (2021) employed LMO3 to drive large-scale chemogenetic excitation of multiple cell types early in development, and then revealed changes in local circuit dynamics after this developmental manipulation using optogenetics in the adult animal *in vivo* [10]. Further, because actuation of BL-OG produces photons, users can track the time window of engagement by measuring this signal. Given the utility of this integrated single actuator approach, we sought to further improve its activation power.

We previously showed that the coupling efficiency between a bioluminescent emitter and optogenetic actuator is readily modifiable, e.g. by pairing different luciferase and opsin variants [11–14]. With bioluminescent activation of an opsin, the magnitude of the total transmembrane current generated (i.e. the efficiency of activity modulation) depends on parameters including the energy output and brightness of the emitter, spectral compatibility (photon emission wavelength with respect to maximum opsin absorption wavelength), photosensitivity of the recipient opsin chromophore, as well as LMO expression level and bioavailability of CTZ.

Here, we sought to improve bioluminescence-driven LMO-mediated modulation of neuronal activity through development of brighter and spectrally matched emitters tuned to optimally drive a sensitive recipient channelrhodopsin. We developed novel luciferases optimized for Förster resonance energy transfer (FRET) when fused to the fluorescent protein mNeonGreen, whose emission is well-tuned to *Volvox* Channelrhodopsin 1 (VChR1). In particular, tethering of the new BL emitter ‘NCS2’ to VChR1 (LMO7, see Table 1) significantly improved the efficiency of activation, more than doubling many of physiological activation metrics, and showing robust control of behavior. This new construct provides the next step in the evolution of this powerful control molecule, displaying superior coupling efficiency together with a retained tracking ability and flexibility for multiple modes of engagement.

**Table 1.** Properties and composition of tested LMOs.

|  | **LMO3** | **LMO5** | **LMO6** | **LMO7** |
| --- | --- | --- | --- | --- |
| **Construct** | sbGluc-VChR1 | Nluc-VChR1 | NCS1-VChR1 | NCS2-VChR1 |
| **Light emitter** | Slowburn *Gaussia* luciferase | Nanoluc | sbAluc1-*d7nm*NeonGreen | mNeonGreend10C-eKL9H |
| **Peak em. $\lambda$ (nm)** | ~470 | ~455 | ~518 | ~518 |
| **Substrate** | Native coelenterazine (N-CTZ) | Coelenterazine-H (H-CTZ)<br>Fluorofurimazine (FFz) | Native coelenterazine (N-CTZ) | Coelenterazine-H (H-CTZ)<br>Fluorofurimazine (FFz) |
| **Actuator** | *Volvox* channelrhodopsin 1 | *Volvox* channelrhodopsin 1 | *Volvox* channelrhodopsin 1 | *Volvox* channelrhodopsin 1 |

## 2 Materials and methods

### 2.1 Animals

All experiments were conducted in accordance with protocols approved by the Institutional Animal Care and Use Committee (IACUC) at respective universities following National Institutes of Health (NIH) guidelines. C57/BL6 (JAX stock #000664) and Swiss Webster (Charles River) mice of both sexes were used and housed 4‒5 animals per cage in a temperature and humidity controlled vivarium on a 12 hour light/dark cycle with ad libitum access to water and food.

### 2.2 Luciferase constructs and characterization of recombinant proteins

Bacterial expression plasmids were constructed using the pNCS backbone, and recombinant protein expression was performed in the *E. coli* strain NEB 10-beta for all experiments as previously described [15, 16]. Directed evolution including error-prone and site-directed mutagenesis and screening for luminescence intensity and wavelength was performed according to methodology described in Lambert et al 2023. Bioluminescence emission spectra were collected using a ClarioStar multimode plate reader (BMG).

### 2.3 Plasmids

The generation of LMO plasmids has been described in detail previously (Berglund et al 2013, Berglund et al 2016). For the new versions of LMOs, the sbGLuc sequence of LMO3 (Berglund et al 2016) was switched out with synthetized mammalian codon-optimized sequences (IDT) of NanoLuc [17], NCS1 and NCS2 (Lambert et al 2023). For the B7 constructs, pcDNA3.1-CMV plasmids were generated featuring sbGLuc or NCS2 tethered to the N-terminus of B7 [18] via the same 15 amino acid linker used for LMOs, with an EYFP tag fused to the C-terminus of B7. For viral vectors, the coding sequence of LMO7 was cloned into an AAV vector downstream of an hSyn promoter. For AAV LMO3, a previously published plasmid was used (Berglund et al 2016; Addgene #114099).

### 2.4 Virus

AAVs carrying LMOs were produced by transfecting subconfluent HEK293FT cells per 10-cm culture dish with 24 μg of the helper plasmid pAd delta F6, 20 μg of the serotype plasmid AAV2/9, and 12 μg of the LMO plasmid using Lipofectamine 2000. After 72 h, the supernatant was harvested from culture plates and filtered at 0.45 μm. Virus was purified from cells and supernatant according to previously described methodology [19], but without the partitioning step in the aqueous two-phase system. Virus was dialyzed against PBS (w/o Ca, Mg) overnight at 4 °C, using FLOAT-A-LYZER G2, MWCO 50 kDa, followed by concentration in Amicon Ultra-0.5 mL Centrifugal Filters. Viral titers were determined by quantitative PCR for the woodchuck hepatitis post-transcriptional regulatory element. Preparations with titers around 1 × 10^13^ vg/mL were used in the study.

### 2.5 Cell culture

HEK293 cells were grown in Dulbecco’s Modified Eagle Medium (DMEM, ThermoFisher) containing 10% fetal calf serum and 0.5% penicillin/streptomycin. Cells were cultured in 6-well plates at 37 °C in a 5% CO2 atmosphere and split every 72 hours by trypsinization (0.25%, Gibco, ThermoFisher). Stable HEK293 lines were generated by lipofection (Lipofectamine 2000, ThermoFisher) of ∼80% confluent cells with 2 µg of linearized expression plasmid overnight according to the manufacturer’s instructions. Cells expressing EYFP and mNeonGreen (LMO3 and LMO5 versus LMO6 and LMO7, respectively) were purified with puromycin titration (1-10 µg/mL, SigmaAldrich) and frozen at -80°C. The percentage of cells showing stable expression ranged from 30‒60% cells in mixed populations.

### 2.6 RNA isolation, cDNA synthesis and RT-qPCR

RNA was extracted from individual wells of 24-well culture plates using a RNeasy Plus Mini Kit (Qiagen) per the manufacturer’s instructions. Culture medium was removed from wells and cells were lysed on the plate in situ. The lysate was homogenized using a QIAshredder spin column, and the homogenized lysate was further purified with a gDNA Eliminator spin column and an on-column DNase digestion. Concentration was determined with a Nanodrop (A260/A280=1.8-2.2; ThermoFisher) and integrity of RNA was assessed in a Bioanalyzer (Agilent). RNA (100ng/sample) was reverse transcribed to cDNA using the High-Capacity RNA to cDNA kit (ThermoFisher) per the manufacturer’s instructions. The generated cDNA (5 ng) was used as a template for RT-qPCR, which was performed with the Applied Biosystems Step One Plus using Power SYBR Green PCR Master Mix (Applied Biosystems, ThermoFisher 4367659). The forward and reverse primers for VChR1 in LMO constructs were 5’-CGGATGGGAGGAGGTTTACG-3’ and 5’-AACTACGCCGTTCCCTGAAG-3’ and for EYFP in B7 constructs were 5’-TAAACGGCCACAAGTTCAGCGTGT-3’ and 5’-ATGTTGCCGTCCTCCTTGAAGTCGA-3’.

### 2.7 IVIS imaging

Luminescence was measured using an IVIS Lumina LT in vivo bioluminescence and fluorescence imaging system (Perkin Elmer). The 4-well plate was moved from the incubator to the imaging room. Medium was removed from each of the 4 wells and replaced with the CTZ-containing medium. Plates were immediately imaged (1 second, no filters, medium binning). Images were displayed as a pseudo-color photon count image. Regions of interest (ROIs) were defined using an automatic intensity contour procedure to identify bioluminescent signals with intensities significantly greater than background. ROIs were measured for each image to obtain values for the Average Radiance [p/s/cm²/sr] for each individual well.

### 2.8 HEK cell electrophysiology

Stable HEK293 cell lines were seeded at low density on 15 mm poly-D-lysine coated coverslips (Neuvitro) in 12-well plates and incubated for 48-72 hours pending electrophysiological analysis. A coverslip was transferred to a recording chamber (RC26-GLP, Warner Instruments) on the stage of an upright microscope (BX51WI, Olympus) and perfused with ACSF (1‒1.5 mL/min) containing (in mM): 150 NaCl, 3 KCl, 10 HEPES, 2 CaCl2, 2 MgCl2 and 20 D-glucose (pH 7.4, ∼300 mOsm/kg) at a temperature of 35 ± 1°C. The intracellular solution contained (in mM): 130 K-gluconate, 8 KCl, 15 HEPES, 5 Na2-phosphocreatine, 4 Mg-ATP, 0.3 Na-GTP (pH 7.25, ∼300 mOsm/Kg). Borosilicate glass micropipettes were manufactured using a vertical puller (PC-100, Narishige) and had resistances from 4‒6 MΩ. Cells expressing EYFP or mNeonGreen were visually targeted in clusters of 5‒15 cells using epifluorescence. VChR1 was excited through the objective (LUMPLFLN40XW, NA 0.8, Olympus) using a metal halide light source (130 W, U-HGLGPS, Olympus) together with filter cubes for blue (Ex/Em: 480/530 nm, U-MNIBA3, Olympus) and green (Ex/Em: 540/600, U-MWIGA3, Olympus) excitation. An electronic shutter (Lambda SC, Sutter Instruments) was used to control light delivery to cells. Photocurrents were evoked by 1 second exposure to blue or green light, respectively, according to peak bioluminescent emission of the evaluated construct (see Table 1). At maximum light intensity the irradiance was 30.8 and 50 mW/cm^2^, respectively (photocurrent amplitudes saturated at lower irradiance for both wavelengths, Fig. S2).

Whole-cell voltage clamp recordings were performed at -60 mV using a Multiclamp 700b amplifier and Digidata 1440 digitizer together with the pCLAMP 10 recording software (Molecular Devices). Data was sampled at 10 kHz, filtered at 3 kHz and analyzed in Igor Pro (WaveMetrics). After break-in, the photocurrent response of the cell was determined using a gap free protocol followed by application of CTZ (Nanolight Technology, #301 and 303) through the perfusion. A fresh stock of CTZ (50 mM in DMSO) was aliquoted at 2 mM and diluted to 250 µM in 1 mL ACSF. Fresh CTZ aliquots were quickly thawed and immediately introduced into the perfusion (final concentration in bath ∼100 µM) for each recording. VChR1 photocurrent-voltage plots were acquired in voltage steps of 10 mV (-80 to +80 mV) during 1 second windows of blue and green light illumination separated by 10 second windows of no light stimulation. The series resistance (Rs) was monitored throughout and recordings were discarded if fluctuations exceeded 25% from start to end. The liquid junction potential was not corrected for.

### 2.9 Brain slice electrophysiology

Acute brain slices were prepared from 3‒6 week-old Swiss Webster mice that received intracortical injection of AAV9-hSyn-LMO virus at postnatal day 2. Mice were deeply anaesthetized via inhalation of isoflurane (Abbott), decapitated, and dissected. The cerebrum was quickly isolated and placed in an ice-cold slicing ACSF containing (in mM): 92 NMDG, 2.5 KCl, 0.5 CaCl2, 10 MgSO4, 30 NaHCO3, 1.25 NaH2PO4, 20 HEPES, 2 Thiourea, 5 Na-ascorbate, 3 Na-pyruvate and 25 D-glucose (310 mOsm/kg, pH 7.3‒7.4) [20]. Coronal sections (300 μm) were prepared from the fronto-parietal cortex of both hemispheres using a vibratome (VT1000s, Leica) and transferred to a chamber containing slicing ACSF at 34 ± 1 °C. Following a NaCl spike-in protocol [20], slices were transferred to a storage solution containing (in mM): 92 NaCl, 2.5 KCl, 2 CaCl2, 2 MgSO4, 30 NaHCO3, 1.25 NaH2PO4, 20 HEPES, 2 Thiourea, 5 Na-ascorbate, 3 Na-pyruvate and 25 D-glucose (310 mOsm/kg, pH 7.3‒7.4) and maintained at RT for up to 6 hrs. After minimum 30 minutes a slice was transferred to a recording chamber mounted on an upright microscope (BX51WI, Olympus) and perfused with ACSF containing (in mM): 121 NaCl, 2.8 KCl, 1 NaH2PO4, 26 NaHCO3, 2 CaCl2, 2 MgCl2 and 15 D-glucose (310 mOsm/kg, pH 7.3‒7.4) at a rate of ∼6 mL/min. All solutions were bubbled with a gas mixture of 95% O2 and 5% CO2. The recording ACSF was supplemented with D-AP5 (50 µM, Tocris), CNQX (15 µM, Tocris) and picrotoxin (100 µM, SigmaAldrich) to block fast glutamatergic and GABAergic synaptic transmission. Whole-cell patch clamp recordings were performed using a Multiclamp 700b amplifier and Digidata 1440 digitizer together with the pCLAMP 10 recording software (Molecular Devices). Borosilicate glass micropipettes were obtained using a PC-100 puller (Narishige) and had resistances of 3‒5 MΩ. Pipettes were filled with intracellular solution containing (in mM): 130 K-gluconate, 10 KCl, 15 HEPES, 5 Na2-phosphocreatine, 4 Mg-ATP, 0.3 Na-GTP and 0.03 Alexa Fluor 594 (ThermoFisher) (310 mOsm/kg, pH 7.3). L5 pyramidal neurons were visually targeted using epifluorescence microscopy together with a CMOS camera (ORCAFusion, Hamamatsu). A current/membrane potential response to blue light was confirmed for each cell prior to recording. Excitation light (480 nm, 1 second, 30.8 mW/cm^2^ or 540 nm, 1 second, 50 mW/cm^2^) was delivered through a 40x water immersion objective using a 130 W metal halide light source (U-HGLGPS, Olympus) together with U-MNIBA3 and U-MWIGA3 filter cubes (Olympus). An electronic shutter (Lambda SC, Sutter Instruments) was used to control the time window of illumination. CTZ was prepared and administered as in HEK cell recordings except at a higher final bath concentration of 300 µM. The effect of CTZ on depolarization and spontaneous firing was examined in continuous recordings from a starting Vm of approximately -70 mV. Vehicle consisted of an equal amount DMSO diluted in ACSF. In LMO7-expressing cells, the effect of CTZ on neural input-output (f-I) was quantified using depolarizing square current injections (800 ms, 0‒300 pA., 25 pA increments) from -70 mV. Bioluminescence was simultaneously recorded (0.2 Hz, 4x4 binning) using an ORCA Fusion CMOS camera (Hamamatsu) in a few cells (LMO3; n = 2, LMO7; n = 3). Data was sampled at 10 kHz, filtered at 3 kHz and analyzed in Igor Pro (WaveMetrics). Rs values were ≤15 MΩ for all cells included in analysis after break-in and were not allowed to fluctuate more than 25% from start to end of the recording. The liquid junction potential was not corrected for. Brain slices containing neurons filled with Alexa Fluor were fixed post hoc in 4% paraformaldehyde (in PBS, pH 7.3‒7.4) overnight at 4°C. Prior to imaging, slices were washed in PBS and cleared using the ScaleSQ(5) protocol [21]. Slices were mounted on slides with ProLong Gold (ThermoFisher), coverslipped and imaged on an Axio Imager M2 microscope (Zeiss) using 10x, 20x and 40x air objectives.

### 2.10 In vivo electrophysiology and bioluminescence imaging

Electrophysiological data was acquired using an Open Ephys acquisition board (http://www.open-ephys.org/) connected via an SPI interface cable (Intan) to a 32-channel headstage (Intan). A 32-channel laminar probe (Neuronexus, A1x32-Poly2-5mm-50s-177) was connected to the headstage. The iridium electrode contacts on the probe covered a linear length of 790 μm and were arranged into two columns of 16 contacts spaced 50 μm apart. The data were acquired using the Open Ephys GUI software at a sampling rate of 30 kHz and referenced to a supra-dural silver wire inserted over the right frontal cortex. Imaging data were acquired using an Andor Ixon Ultra 888 EMCCD camera fitted with a Navitar Zoom 6000 lens system together with the Andor Solis data acquisition software (Andor Solis 64 bit, v4.32). The field-of-view was centered over the craniotomy and adjusted to encompass the full 3 mm diameter of the craniotomy. Images (512 x 512 pixels, ∼6 μm2/pixel) were acquired continuously at an exposure length of 2 seconds and an electron multiplication gain of 300. To test the optogenetic elements of the constructs, external 530 nm light was delivered by an LED (Mightex FCS-0530-0000) coupled to an optic fiber (105 μm diameter, 0.22 NA). The patch fiber was coupled to a fiber on the neuronexus probe (105 μm diameter, 0.22 NA), which terminated 200 μm above the top contact of the probe.

Experiments were conducted approximately seven weeks following surgery (LMO7: M = 6.98, SD = 1.60, median = 6.71, range = 5.00‒9.71; LMO3: M = 7.18, SD = 1.94, median = 7.14, range = 4.85‒10.14). On the day of the experiment, the animal was anesthetized (1‒2% isoflurane) and a dental drill was used to remove the cement holding the window in place. The window was then lifted off to expose the open craniotomy. For the duration of the experiment the craniotomy was kept covered in saline. The animal remained anesthetized for the entirety of the experiment and was transcardially perfused at its conclusion.

We identified the regions of strongest viral transfection within the craniotomy under epifluorescent 530 nm illumination. For each animal, the center of this region was selected for the location to insert the electrode, avoiding visible blood vessels. The electrode was lowered into the cortex through the intact dura using a micromanipulator (Siskiyou MD7700) at a rate of 1 μm/sec. The electrode was lowered until the uppermost contact on the laminar probe disappeared from view into the brain as viewed under a stereoscope. The probe was then left undisturbed for 30 minutes before beginning recording. Each animal was next tested for the presence of an LED evoked optogenetic response. Fifty trials of two gaussian pulses (peak intensity at fiber tip = 10.7 mW/mm^2^) 200 ms in length with a 200 ms inter-pulse-interval were delivered at an inter-trial-interval of 7 to 10 seconds (pseudorandom square distribution). Multi-unit spiking activity (MUA) was monitored during stimulation. All animals exhibited an increase in MUA with LED stimulation that was readily observable in real-time on a trial to trial basis.

Either water soluble native CTZ (N-CTZ, LMO3 animals, Nanolight CAT#3031) or coelenterazine H (H-CTZ, LMO7 animals, Nanolight CAT#3011) was dissolved in sterile saline (4.55 μg/μL) to yield a final concentration of 11 mM. The solution was loaded into a 100 μl glass syringe (Hamilton #80601) fitted with a ∼1 cm length of 18 gauge plastic tubing. The Hamilton syringe and tubing was placed in a motorized injector (Stoelting Quintessential Stereotaxic Injector, QSI). The tip of the plastic tubing was lowered into the pool of saline over the craniotomy using a micromanipulator until it touched the surface of the skull. The tip of the tubing was further adjusted so that it rested at a distance of ∼7.5 mm from the opening edge of the craniotomy. The CTZ was delivered by infusing 20 μL of the solution into the saline over the open craniotomy at a rate of 200 μL/min. The volume of the saline well within the headpost and surrounding the craniotomy was measured to be 200 μL in volume. The addition of the 20 μL solution to the saline was chosen to yield a 1 mM final concentration in the saline well.

A TTL pulse co-triggered the acquisition of electrophysiology and imaging data and a minimum of 5 minutes of baseline activity was recorded for each animal. The CTZ was then infused into the craniotomy and data were recorded for a minimum of 20 minutes.

### 2.11 Behavioral testing

Three female C57BL/6 mice were used. Several weeks after viral transduction the behavior of mice in an open field chamber was assessed after intraperitoneal (i.p.) injection of vehicle (saline) and fluorofurimazine (FFz; 175 μL, 1.53 µmoles) [22, 23] in two recording sessions on separate days. Immediately after the injection, mice were placed in the open field chamber (clear plexiglass, 10”x12”x15”). RaspberryPi 4 cameras were utilized to capture videos (1280 x 730 at 25 fps) from below with recordings starting 5‒8 minutes after the i.p. injection. Motor behaviors were tracked for 10 minutes. Videos were analyzed for overall turning preferences (ipsi- vs contralateral of the transduced SNr) using DeepLabCut. Number of rotations, grooming bout number and total time spent grooming were manually scored by an experimenter blind to the treatment (vehicle, FFz) for each mouse.

### 2.12 Surgery and viral injections

#### 2.12.1 Brain slice electrophysiology

P2 Swiss Webster pups were cryo-anesthetized on ice for 5 minutes prior to injection. Upon cessation of movement pups were bilaterally injected with 1 µL AAV9-hsyn-LMO3-EYFP (3 females/3 males) or AAV9-hsyn-LMO7 (4 females/3 males) into the fronto-parietal cortex using a borosilicate glass pipette. Injections were performed manually by applying a small positive pressure to the pipette. Prior to, and following injections, pups were kept warm on a heating blanket and placed back with their dam immediately after the procedure.

#### 2.12.2 In vivo electrophysiology

A total of 15 C57BL/6J mice (7 female/8 male) were used. Seven of these animals (3 females/4 males) were injected with the LMO3 viral construct (AAV9-hsyn-LMO3-EYFP; age on day of surgery: M = 26.92 weeks, SD = 4.18, median = 30.14, range = 26.00‒40.43). The remaining eight animals (4 females/4 males) were injected with the LMO7 viral construct (AAV9-hSyn-LMO7; age on day of surgery: M = 26.23 weeks, SD = 4.79, median = 30 , range = 26.28‒40.28). Each animal was anesthetized (1‒2% isoflurane), fitted with a steel headpost, and injected with viral constructs in a 3 mm craniotomy centered over left SI (-1.25 A/P, 3.5 M/L relative to bregma) made with a dental drill. Three injections of the constructs were made at locations equidistantly spaced around the central SI point at depths of 500 μm. Each injection was 400 nL in volume and delivered at a rate of 50 nL/min. Viral injections were performed through a glass pipette fitted in a motorized injector (Stoelting Quintessential Stereotaxic Injector, QSI). Finally a glass window was fitted over the open craniotomy and cemented in place with dental cement (C & B Metabond). Dexamethasone was given intraperitoneally (0.5 mg/kg) to reduce brain edema, and sustained-release meloxicam (Meloxicam SR) was given subcutaneously (4 mg/kg) for pain relief during recovery.

#### 2.12.3 Behavior

Ten-week-old mice were stereotaxically injected into the substantia nigra pars reticulata (SNr) with the LMO7 viral construct (AAV9-hSyn-LMO7) through a unilateral craniotomy (from Bregma: A/P -3.4; M/L 1.25; D/V -4.2). Solution containing virus at a concentration of 1 x 10^13^ vg/mL was injected at a volume of 500‒800 nL and at a rate of 100 nL/min using a NanoFil syringe with a 35-gauge beveled needle.

### 2.13 Data analysis

#### 2.13.1 HEK cell electrophysiology

Photocurrent amplitudes were analyzed at steady state level as the average current amplitude measured during the last 100 ms of illumination to circumvent inconsistencies associated with light-adaptation and the peak current response. The CTZ-induced current amplitude was quantified at the peak plateau level observed within 5 minutes as a ± 10 second average around the peak amplitude. Only recordings where initial holding current was below -100 pA and stable for minimum 2 minutes following light stimulation were used in analysis. The CTZ-induced response was normalized to photocurrent amplitude in each recorded cell to yield the coupling efficiency (CTZ-induced current amplitude / maximum photocurrent amplitude). Only cells displaying photocurrent amplitudes of minimum 500 pA were used for evaluation of coupling efficiency. In non-expressing controls the effect of CTZ on holding current was quantified 5 minutes post wash in and averaged over 20 seconds. Data was analyzed in Clampfit (Molecular devices) and Igor Pro (WaveMetrics).

#### 2.13.2 Brain slice electrophysiology

Cells displaying holding currents less than -150 pA and Rs ≤ 15 MΩ were included in the analysis. CTZ-induced depolarization amplitudes were calculated as a ± 5 second average around a manually identified peak amplitude observed at 50‒80 seconds following substrate delivery to bath. The effect of vehicle on Vm was calculated as a 10 second average 70‒80 ms post substrate addition (no effect on Vm was observed after an additional 60 seconds). In frequency-current (f-I) plots the number of action potentials produced during 800 ms depolarizing square current injection was calculated at baseline (pre), 90 seconds post CTZ delivery to bath (CTZ) and following 5 minutes of wash out with ACSF (post). The rheobase current was defined as the minimum stimulation magnitude required to evoke ≥ 1 action potentials. Action potential threshold was defined as a dV/dt of ≥ 20 mV/ms and measured from the first action potential generated by the depolarizing current protocol. Bioluminescence traces were generated by plotting time-lapse data as mean intensity values from somatic ROIs of LMO-expressing L5 pyramidal neurons in ImageJ, and time-synchronized to membrane potential recordings. In simultaneous bioluminescence and Vm recordings the peak intensity was normalized to the peak depolarization amplitude. All data was analyzed in Igor Pro (WaveMetrics).

#### 2.13.3 In vivo electrophysiology and imaging

Data were not used from one LMO3 mouse due to a bad reference connection resulting in excessive noise in the data and no discernible spiking response to LED stimulation, and one LMO7 mouse due to a hardware malfunction during the infusion of CTZ. Offline analyses and statistical tests of both electrophysiological and imaging data were performed in Matlab R2022a (Mathworks). For each recording, electrodes with RMS values more than three times the interquartile range above the 3rd quartile or less than three times the interquartile range below the 1st quartile of all 32 electrodes were marked as excessively noisy and removed from further analyses. (LMO7 number of bad electrodes: Mean = 2.28 , SD = 0.75 , median = 2 , range = 1‒3; LMO3 number of bad electrodes: Mean = 2.67, SD = 1.21 , median = 2.5, range = 1‒4). The remaining electrodes were re-referenced to the common average reference [24]. To isolate MUA, a bandpass filter (passband: 500 Hz to 5000 Hz, 3rd order Butterworth) was applied to the data in the forwards and backwards directions to avoid phase distortions. Spikes were defined as data points less than -3 times the standard deviation, where the standard deviation was estimated as the median divided by 0.6745 [25]. Each electrode’s spike time series was then convolved with a 50 ms gaussian window (SD = 5 ms) to provide an estimate of the instantaneous firing rate.

In the case of LED stimulation, the MUA data were epoched into trials -2 to 2 seconds relative to stimulus onset. Next we identified responsive electrodes, defined as electrodes in which the lower bootstrapped 99% CI of the poststimulus response (mean MUA activity 0 to 600 ms after LED onset) was greater than the upper bootstrapped 99% CI of the prestimulus 2 second baseline, pooling all the trial data (number of responsive electrodes LMO7: M = 30.85, SD = 0.89, median = 31, range = 29‒32; LMO3: M = 27.67, SD = 4.93, median = 29, range = 18‒31). The data were then averaged across trials and responsive electrodes to produce a single LED evoked waveform for each animal and converted to percent change relative to baseline. LED response values for each animal are reported as the average percent change from baseline in the 0 to 600 ms poststimulus time period. Spike time series were binned at 1 second intervals in the case of CTZ administration. CTZ responsive electrodes were identified as those whose lower 99% bootstrapped CI of the MUA after CTZ delivery was greater than the upper 99% bootstrapped CI of the baseline period baseline (Number CTZ responsive electrodes LMO7: M = 11.28, SD = 4.85, median = 10, range = 4‒16; LMO3: M = 15.17, SD = 8.35, median = 14.5, range = 4‒24). CTZ responsive electrodes were then averaged for each animal and converted to percent change relative to baseline. Among all animals, all electrodes identified as CTZ responsive were also identified as LED responsive. CTZ response values are reported as the maximum increase from baseline in a 60 second moving average across the entire post CTZ time period.

For the imaging data, each image was binned offline by taking the median value in an 8x8 pixel bin, resulting in a new image size of 64 x 64. For each animal, a circular region with a diameter of 8 pixels (∼376 μm) was placed in the region directly adjacent to the electrode shank and in front of the surface with the exposed electrode contacts. The mean of these pixels was computed for each image to yield a time series of bioluminescent light output. Each animal’s bioluminescence time series was then converted to percent change relative to baseline. Bioluminescence response values for each animal were calculated as maximum increase from baseline in a 60 second moving average across the entire post CTZ time period. Unless otherwise stated all significance testing was performed using independent samples t-tests (Matlab ttest2).

#### 2.13.4 Behavior

DeepLabCut 2.1, run on a Lenovo Thinkstation P620 with a NVIDIA Quattro RTX 5000 graphics card, was utilized for markerless quantification of body orientation by tracking the nose, four paws, and the base of the tail [26, 27]. Three hundred frames were selected for labeling and used to train the network with a ResNet101 based neural network, utilizing 95% of the labeled frames with default parameters for a single training session. The network was trained for 200,000 iterations until the loss reached a plateau. By validating with three shuffles, the test error with a p-cutoff value of 0.9 was 4.39 pixels. We used a p-cutoff of 0.9 to condition the X,Y coordinates for further analysis. Segments of recordings with a likelihood of <0.9 were excluded from analysis. Custom scripts were utilized to quantify cumulative instantaneous body angle after vehicle and FFz i.p. injections. The body angle was calculated based on the tail base to nose vector relative to the x-axis.

Statistical significance was defined as P-values <0.05 and evaluated using paired or unpaired t-tests, or Kruskal-Wallis tests, after testing for normality of data. SPSS (IBM) and Matlab (Mathworks) softwares were used for statistical analysis. Data is reported as mean ± SEM unless otherwise specified.

## 3 Results

### 3.1 Generation of FRET-based LMOs

As strategy to generate more efficient LMOs, we focused on new light-emitters displaying brighter BL emission and/or improved spectral compatibility with the *Volvox* channelrhodopsin 1 (VChR1) chromophore. To this end, we engineered highly efficient FRET pairs, using mNeonGreen as the acceptor, by (i) designing a novel bright copepod- derived luciferase and (ii) evolving an *Oplophorus gracilirostris* luciferase (OLuc) [28] variant with improved performance compared to NanoLuc [17].

To generate an improved copepod-type luciferase, we designed several variants by introducing subsets of mutations identified by other groups for *Gaussia princeps* luciferase (Gluc) [29, 30] into consensus-derived “artificial” luciferase (Aluc) sequences [31]. The clone yielding the highest luminescent signal with CTZ when produced in *E. coli* was chosen for full characterization and named “sbALuc1” to reflect its major derivation from “sbGluc” mutations paired with an Aluc. We next optimized a fusion construct of sbALuc1 with mNeonGreen following a similar approach to that taken to optimize the mVenus-RLuc8 fusion Yellow NanoLantern (YNL) [32]. Ultimately, this approach generated an optimized FRET construct code-named “NCS1” which was used as light emitter to generate LMO6 (Fig. 1A-C, and Table 1).

**Fig. 1.**
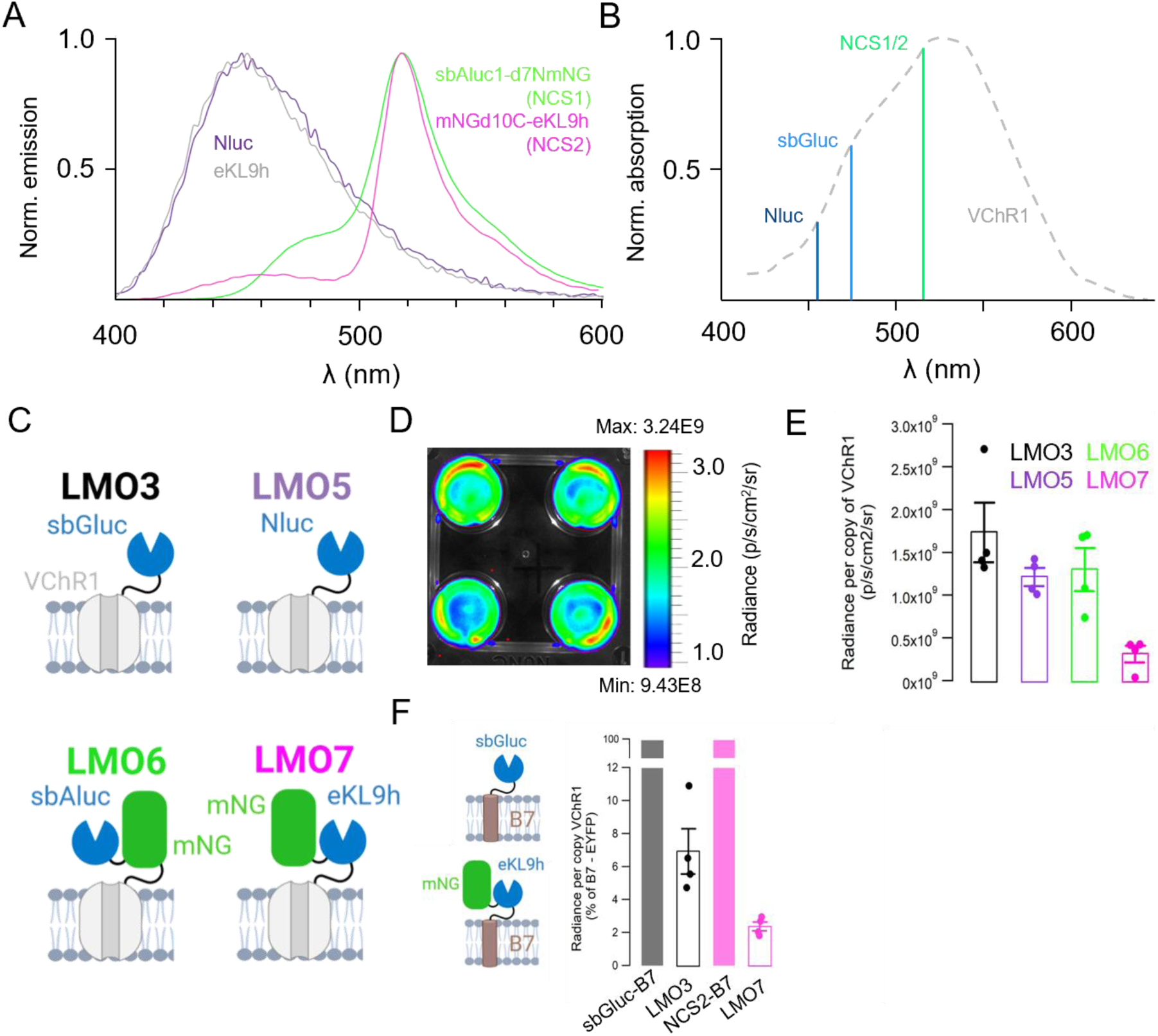
Novel luciferase FRET pair characteristics. **(A)** Bioluminescence emission spectra from the luciferases NanoLuc (Nluc) and eKL9h compared to the mNeonGreen-luciferase fusions NCS1 and NCS2. (**B**) Action spectrum of VChR1 compared with the peak emission wavelengths of NLuc, sbGluc, and NCS1/NCS2 illustrating the better alignment achievable using FRET. (**C**) Schematic of the LMOs compared for bioluminescence emission in HEK cell cultures. (**D**) Example image of bioluminescence recorded from HEK cell cultures in 4-well plates using IVIS imaging. (**E**) Summary bar graph showing the average radiance per copy of VChR1 recorded in the four groups. (**F**) Summary bar graph showing the average radiance of LMO3 and LMO7 relative to the radiance recorded from B7-tethered constructs (left).

We also generated an OLuc variant with favourable properties for fusion constructs as part of a larger effort to generate bright luminescent probes (described in greater detail in Lambert et al., 2023). Briefly, we performed multiple rounds of directed evolution consisting of alternating error-prone and site-directed mutagenesis libraries in *E. coli*. For each round, variants were selected based on higher bioluminescent emission and solubility. We subsequently introduced a subset of mutations found in NanoLuc [17] and NanoBit [33] to generate the clone “eKL9h,” which was fused to mNeonGreen using the same end truncations and linker sequence as used for the mNeonGreen-NanoLuc pair GeNL [34]. A final module (‘NCS2’) was obtained and used as light emitter to generate LMO7 (Fig. 1A-C, Table 1). Notably, NCS2 is fully soluble when expressed in *E. coli* while GeNL, which differs only in the luciferase domain (NanoLuc), is largely insoluble [35]. We therefore chose NCS2 over GeNL as the bioluminescent component of our new LMO construct.

Because the mNeonGreen FRET acceptor produces bioluminescence emission peaking ∼518 nm for NCS1 and NCS2, we expected both to activate VChR1 more efficiently than blue-emitting luciferases (Fig. 1B). To test the ability of NCS1 and NCS2 to drive opsin activation, we generated new LMOs by fusing them to the N-terminus of VChR1 alongside NanoLuc (Nluc) for comparison (Fig. 1C and see Table 1 for information on tested LMOs). We first examined the light output from LMOs in transiently transfected HEK293 cells by IVIS imaging to obtain values for the average radiance [p/s/cm²/sr] for each individual well (see Fig. 1D for an example). To baseline light emission to single molecules, RNA was prepared from duplicate plates, reverse transcribed and used for qPCRs to determine the copy number of VChR1 molecules per well. As the light emitters are tethered to the opsin, this allows determining the radiance per copy of LMO. Figure 1E shows the average radiance [p/s/cm²/sr] recorded for each LMO. Interestingly, radiance was lowest from LMO7, leading us to speculate that FRET from the light emitter (here the mNeonGreen-luciferase fusion protein) to the nearby VChR1 chromophore is more efficient for NCS2 than for the other light emitters. To explore this further, we tethered sbGLuc and NCS2 to the extracellular side of the B7 transmembrane sequence from the mouse CD80 antigen and compared their light emission when tethered to an opsin versus a transmembrane sequence. With light emission per copy number set at 100% for the B7 tethered constructs, radiance per copy decreased for both opsin tethered light emitters, with a larger decrease for LMO7 versus LMO3 (Fig. 1F). This is consistent with a loss of photon emission to direct energy transfer being more efficient for LMO7 than for LMO3.

### 3.2 FRET-based LMOs generate superior bioluminescence-driven inward currents in HEK293 cells

To compare the ability of new LMOs to drive VChR1-mediated inward currents, we generated stable expression HEK293 lines for LMO3, 5, 6 and 7, and used whole-cell patch clamp recordings to determine the coupling efficiency of each construct (i.e. the ratio of luciferin-induced versus maximum LED-induced photocurrent amplitude response). Expressing cells were identified with epifluorescence and voltage-clamped at -60 mV. At testing outset, brief light stimulation (blue or green, 1 second) was applied through the 40x water immersion objective to drive a photocurrent response. Depending on the construct, either native or H-CTZ (100 µM) was then introduced via an ACSF perfusion to cells during continuous recording (Fig. 2). In non-expressing (control) HEK293 cells, neither substrate created a membrane current (Fig. 2B, E and I). However, in LMO-expressing cells robust CTZ-driven inward currents were observed for all four constructs (Fig. 2C, D, F, G). These inward currents developed gradually over 3-4 minutes to reach plateau levels within 5 minutes of substrate infusion to the bath (Fig. 2J). We found that the four LMOs generated varying CTZ-driven inward current amplitudes on average (LMO3: -131.1 ± 32.7 pA, n = 9; LMO5: -87.1 ± 33.3 pA, LMO6: -251.4 ± 44.4 pA, n = 10; LMO7: -659.1 ± 103.9 pA, n = 9; Fig. 2I, J, Kruskal-Wallis test, p <0.001; pairwise Dunn’s comparison test with LMO3: p = 0.41 (LMO5), p = 0.19 (LMO6), p = 0.003 (LMO7), whereas photocurrent amplitudes and densities were very similar between groups (Fig. S1). As a result, the coupling efficiency also varied between LMOs (LMO3: 13.4 ± 3.4%, n = 9; LMO5: 7.2 ± 1.7%, LMO6: 24.8 ± 3.7%, n = 10; LMO7: 63.7 ± 7.2%, n = 9; Fig. 2K, Kruskal-Wallis test, p <0.001; pairwise Dunn’s comparison test with LMO3: p = 0.33 (LMO5), p = 0.06 (LMO6), p = 0.0002 (LMO7). LMO7 showed the largest average CTZ-driven current amplitude and coupling efficiency, more than 4-fold greater than the prior standard LMO3.

**Fig. 2.**
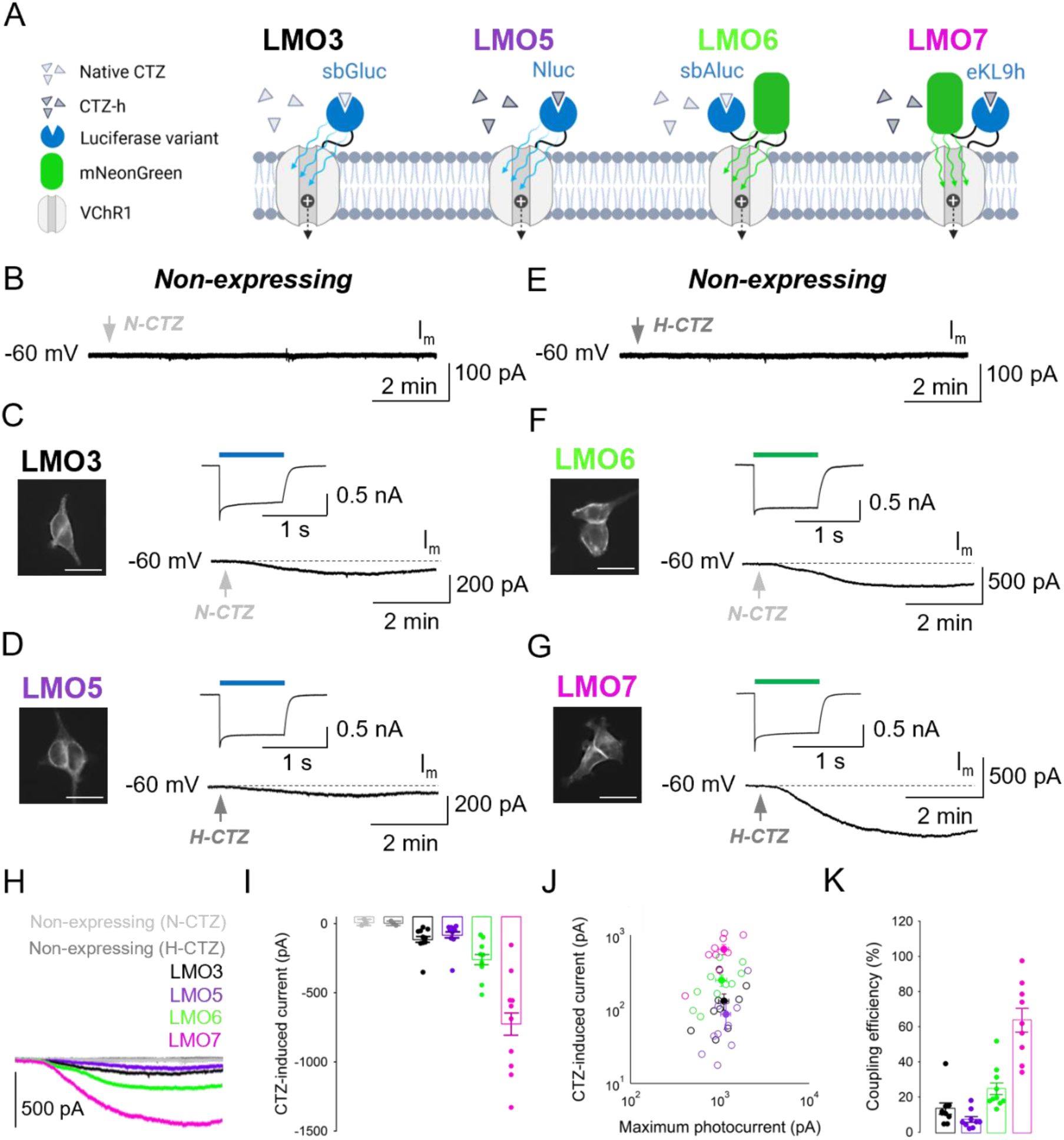
Evaluation of the coupling efficiency of new LMOs in HEK cells. (**A**) Schematic of the four LMOs, with respective CTZ substrates, tested for coupling efficiency. (**B**) Effect of native coelenterazine (N-CTZ) on membrane current (Im) in a non-expressing HEK cell. (**C**) Left, EYFP fluorescence imaged in a stable HEK line expressing LMO3. Right, photocurrent response evoked by 480 nm light (1 second) and example trace showing the representative inward current amplitude recorded 5 minutes post N-CTZ application. (**D**) Left, EYFP fluorescence imaged in a stable HEK line expressing LMO5. Right, photocurrent response evoked by 480 nm light (1 second) and example trace showing the representative inward current amplitude recorded 5 minutes post H-CTZ application. (**E**) Effect of coelenterazine H (H-CTZ) on Im in a non-expressing HEK cell. (**F**) Left, mNeonGreen fluorescence imaged in a stable HEK line expressing LMO6. Right, photocurrent response evoked by 540 nm light (1 second) and example trace showing the representative inward current amplitude recorded 5 minutes post N-CTZ application. (**G**) Left, mNeonGreen fluorescence imaged in a stable HEK line expressing LMO7. Photocurrent evoked by 540 nm light for 1 second. Right, photocurrent response evoked by 540 nm light (1 second) and example trace showing the representative inward current amplitude recorded 5 minutes post H-CTZ application. (**H**) Example trace comparison of CTZ-induced inward current amplitude across stable HEK lines as indicated. (**I**) Summary bar graph of average CTZ-induced current amplitude generated with each construct. (**J**) Summary graph showing correlation between CTZ-induced current and maximum photocurrent in all cells (**K**) Summary bar graph showing the coupling efficiency (CTZ-induced current amplitude / maximum photocurrent amplitude) achieved with each construct.

The excitation wavelength used to photoactivate VChR1 was matched to the peak emission wavelength of the BL emitter (see Table 1), blue light (480 nm) excitation for LMO3 and 5, and green light (540 nm) for LMO6 and 7. To confirm that this protocol allowed for side-by-side comparison of LMOs, we recorded photocurrent responses to blue and green light from the same cell in a subset of LMO3 and LMO7 expressing cells (Fig. S2). At the irradiance level used in the above set of patch clamp experiments (480 nm: 31 mW/cm^2^; 540 nm; 50 mW/cm^2^), evoked photocurrents showed no significant difference in voltage dependence or reversal potential (Fig. S2A, B) and were of similar amplitude (Fig. S2C, D) in response to blue and green light stimulation.

### 3.3 LMO7 generates stronger depolarization in neocortical neurons ex vivo

We next tested the ability of LMO7 to drive membrane depolarization in neurons relative to LMO3. Postnatal day 2 (P2) mouse pups were bilaterally injected with AAV encoding LMO3 or LMO7 under the human synapsin promoter (hSyn) in fronto-parietal cortex, and coronal slices were prepared at 3-6 weeks. For both LMOs, expression was observed primarily in layer 5, as verified with 480 nm epifluorescence prior to experiments and confirmed with post hoc imaging (Fig. S3). LMO-expressing cells were randomly targeted for whole cell recordings and showed predominantly pyramidal morphological and electrophysiological properties (Fig. 3A, D). To verify opsin expression, cells were exposed to a 1 second pulse of 480 nm (31 mW/cm2) and 540 nm (50 mW/cm2) light after break-in to record the photocurrent amplitude and confirm light-evoked firing responses (Fig. 3A, D, right). Following a stable baseline recording period at resting membrane potential, CTZ substrates were applied at equal concentration (∼300 µM final concentration in bath) through the ACSF perfusion. Upon CTZ delivery, bioluminescent emission typically peaked during the first 60 seconds after onset, followed by a reversible depolarization of the membrane potential (Fig. 3B, E) of variable magnitude (Fig. 3H). We found that LMO7-expressing neurons on average produced a ∼2-fold greater depolarization relative to LMO3 (10.5 ± 1.8 mV vs 5 ± 1 mV, Fig. 3G, H, p = 0.019, unpaired t-test). In control experiments, vehicle application had no effect on the membrane potential (Fig. 3B, E, H), and average photocurrent amplitudes were similar between cell populations (Fig. S4, LMO3: -1191.5 ± 186 pA, LMO7: -1112,9 ± 166 pA, p = 0.76, unpaired t-test). In a small number of cells (LMO3: 1/10, LMO7: 3/11) CTZ application produced suprathreshold depolarization (Fig. 3C, F). To verify that LMO7 increases the firing output of neurons, we also recorded the frequency-current (f-I) curve in a subset of neurons (n = 7) pre, during and post CTZ application (Fig. 3I-L). We found that CTZ application significantly lowered the rheobase current (100 ± 7.7 pA vs 64.3 ± 5.1 pA, Fig. 3K, n = 7, p = 0.0004, paired t-test) and increased the maximum number of action potentials produced (18 ± 1.8 vs 21 ± 2.2, Fig. 3L, n = 7, p = 0.01, paired t-test), causing a leftward shift of the f-I curve (Fig. 3J). Statistically significant firing responses were observed in the 50‒175 pA range (p < 0.05, paired t-test).

**Fig. 3.**
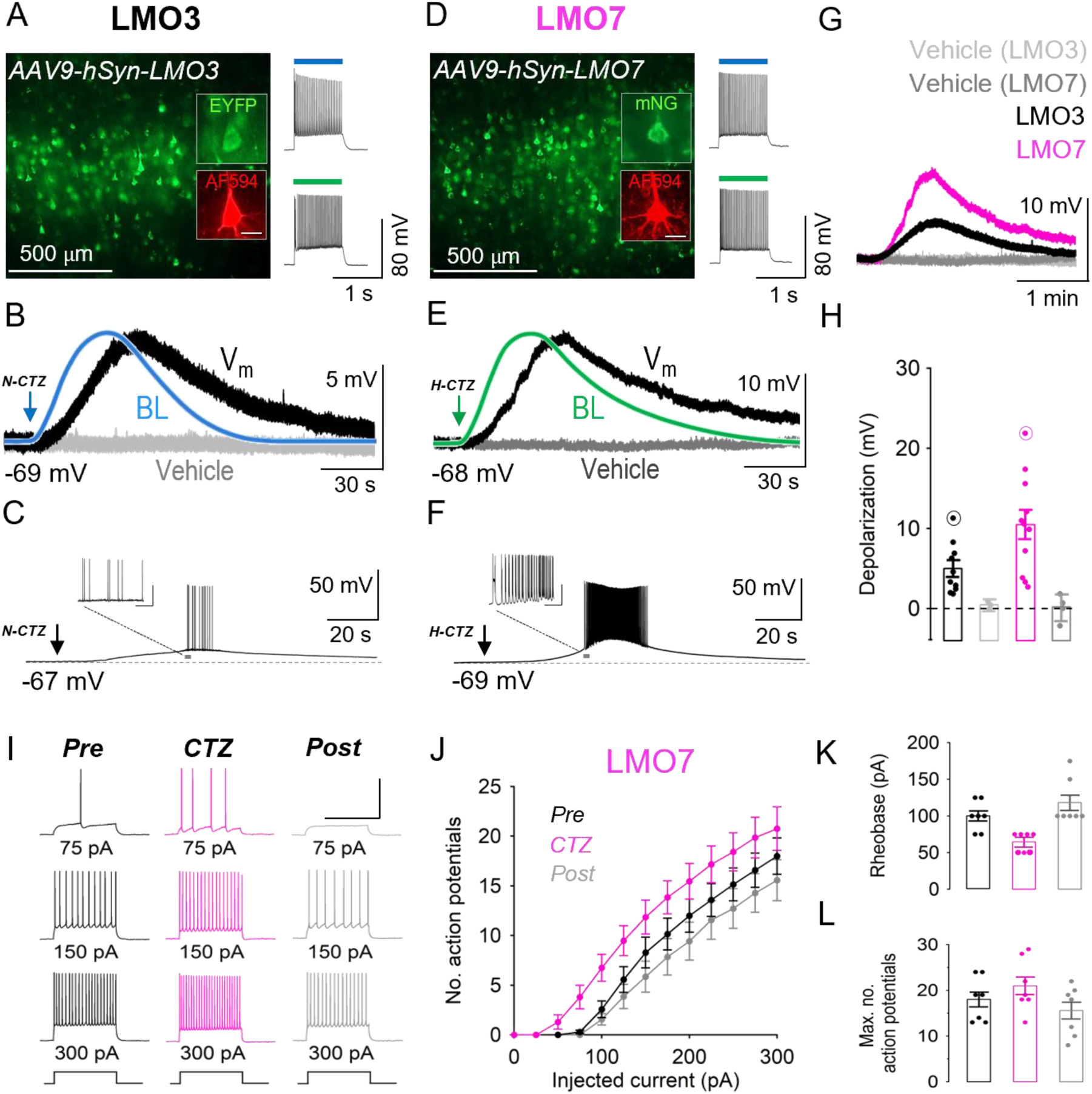
Comparison of LMO3 and LMO7 in acute mouse brain slices. (**A**) LMO3 expression in L5 neocortical neurons from fronto-parietal coronal slices (insets show a pipette-filled pyramidal cell visualized with AlexaFluor 594 and the EYFP tag post hoc). Right, firing response of the cell in response to 480 nm and 540 nm excitation. (**B**) Example trace showing the membrane potential response of the cell in *A* to N-CTZ (black trace, Vm) normalized to the recorded peak bioluminescence (blue, BL) and relative to control (vehicle, light grey trace). (**C**) Example trace showing a suprathreshold Vm response recorded from a LMO3-expressing L5 pyramidal neuron to CTZ. (**D**) LMO7 expression in L5 neocortical neurons from fronto-parietal coronal slices (insets show a pipette-filled pyramidal cell visualized with AlexaFluor 594 and mNeonGreen post hoc). Right, firing response of the cell in response to 480 nm and 540 nm excitation. (**E**) Example trace showing the membrane potential response of the cell in *D* to H-CTZ (black trace, Vm) normalized to the recorded peak bioluminescence (green, BL) and relative to control (vehicle, dark grey trace). (**F**) Example trace showing a suprathreshold Vm response recorded from a LMO7-expressing L5 pyramidal neuron to CTZ. (**G**) Representative example traces of CTZ-induced depolarization amplitude in LMO3 and LMO7-expressing cells relative to controls. (**H**) Summary graph showing the average CTZ-induced depolarization (circled data points represent recordings in *C* and *F*). (**I-L**) Effect of H-CTZ on the input-output function of LMO7-expressing L5 pyramidal neurons. (**I**) Example traces showing effect of CTZ on firing rate at indicated levels of square current injection. (**J**) Summary graph showing the frequency-current response of LMO7-expressing cells pre, during- and post CTZ application. (**K, L**) Summary bar graphs showing effect of CTZ on rheobase (**K**) and maximum number of action potentials (**L**) in LMO7-expressing L5 pyramidal neurons. Scale bars (A, D): 40 µm.

### 3.4 LMO7 elicits larger bioluminescence-driven MUA responses in mouse somatosensory cortex

We proceeded to test the impact of LMO7 activation *in vivo* in Primary Somatosensory (SI) Neocortex of adult wild type mice in comparison to LMO3. Seven weeks post viral injection (AAV9-hSyn-LMO3 or LMO7) all animals showed clear MUA responses to LED stimulation (Fig. 4, panel C, percent change from baseline, LMO7: Mean = 729.21, SD = 353.48, median = 844.07, range = 322.9‒1197.9; LMO3: Mean = 1233.2, SD = 1130.9, median = 1047.7, range = 164‒3285). Although the LMO3 group demonstrated greater responses to the LED on average than the LMO7 group, this difference did not reach statistical significance (t (11) = -1.12, p = 0.28). All animals showed bioluminescence after CTZ delivery (Fig. 4, panel B; percent change from baseline, LMO7: Mean = 273.3, SD = 216.2, median = 237, range = 25.93‒572.2; LMO3: Mean = 739.7, SD = 590.1, median = 741.3, range = 58.74‒1517.8). Bioluminescent output was greater on average in the LMO3 group than the LMO7 group, and this difference showed a trend to significance (paired t-test: t (11) = -1.96, p = 0.076).

**Fig. 4.**
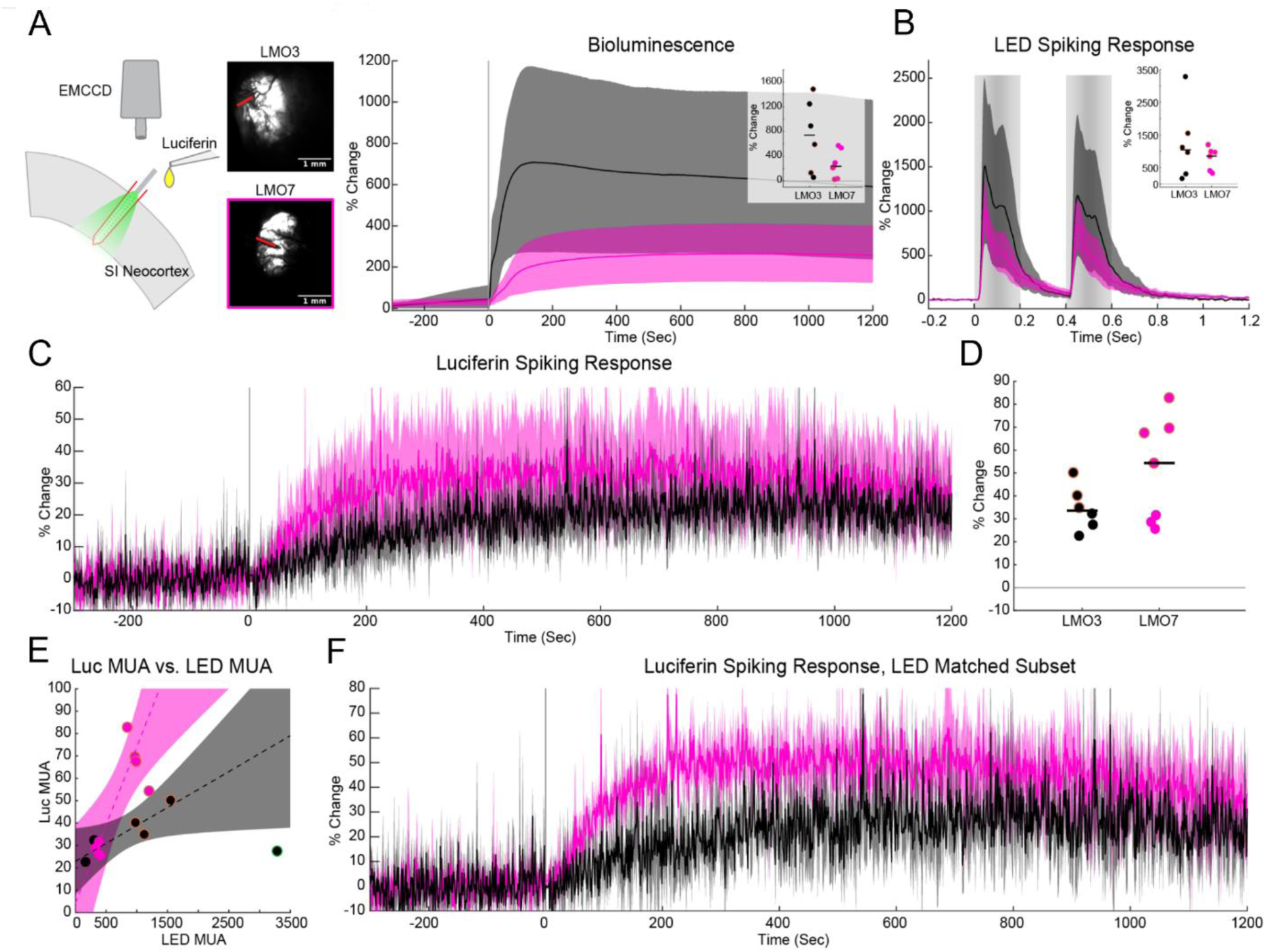
In vivo comparison of LMO3 and LMO7 (black and magenta throughout, respectively*. (***A**) Schematic of the *in vivo* experiment. The laminar probe was inserted perpendicularly to the primary somatosensory cortex. A fiber attached to the electrode delivered light stimuli. The luciferin (native or H-CTZ) was infused over the top of the cortex, and bioluminescent light output was recorded using an EMCCD camera. Example images of bioluminescence for each group are depicted to the right of the schematic. The red lines indicate the location of the electrode in each instance. (**B**) Average bioluminescent light output for the two groups expressed as the percent change from baseline. Shaded error bars indicate the bootstrapped 95% confidence interval. Inset: Dots are individual animal’s peak bioluminescent output, horizontal black bars indicate the median for each group. (**C**) Average MUA response to the external LED stimulus for each group. Shaded gray bars indicate the relative intensity of the LED stimulus through time. (**D**) The average MUA response to luciferin delivery for the two groups. The onset of the 60 second luciferin infusion began at time 0. (**E**) The linear fit of animals’ LED evoked MUA to luciferin evoked MUA. Dashed lines are the linear fits for the two groups and shaded error bars are the 95% confidence intervals of the fitted lines. (**F**). The average luciferin evoked MUA activity of the two LED evoked activity matched animal subsets.

Given that VChR1 is the opsin in both constructs, the trend to higher LED and MUA responses in the LMO3 mouse indicated higher levels of expression. In support of this conclusion, across the two groups and within each group, there was a strong linear relationship between CTZ driven MUA and LED driven MUA (Fig. 4, panel E). A robust linear fit of the CTZ MUA to the LED driven MUA, collapsing across the two groups was significant (R^2 = 0.596, F (1,11) = 15.3, p = 0.002), an effect also observed within each group (LMO7: F (1,5) = 16.8, p = 0.009, R^2 = 0.775; LMO3: F(1,3) = 11.4, p = 0.04, R^2 = 0.79).

To control for differences in expression, we chose a matched subset of animals (LMO3: n = 3, LMO7: n = 4) in which the LED driven responses clustered closely about the median LED responses of the two groups (LMO7 LED subset response: Mean = 999.5, SD = 146.8, median = 978, range = 844.1‒1197.9; LMO3 LED subset response: Mean = 1215.1, SD = 298.7, median = 1119.1, range = 976.2‒1549.9, Fig. 4; orange outlined dots throughout).

When these matched subgroups were compared, LMO7 drove a 64.2% greater CTZ driven MUA response (Mean = 68.6 Hz, SD = 11.7, median = 68.6, range = 54.3‒82.8) than the LMO3 subgroup (Mean = 41.8 Hz, SD = 7.8, median = 40.2, range = 34.9‒50.2; t (5) = 3.41, p = 0.019).

### 3.5 LMO7 expression in the substantia nigra drives robust motor behavior in mice

To test its utility for studying behavior, we measured motor turning in freely moving mice that unilaterally expressed LMO7 within the substantia nigra pars reticulata (SNr; Fig. 5A and Fig. S5). While the native *Oplophorus* luciferase (OLuc) uses coelenterazine as its luciferin, engineered luciferases derived from OLuc emit more light with the coelenterazine analogue furimazine [17]. In addition, furimazine has been chemically modified to provide fluorinated analogs, such as fluorofurimazine (FFz), with improved pharmacokinetics and rapid onset of peak light emission *in vivo* [22, 23]. FFz has been reported to reach maximal photon emission in mouse brain 5 minutes after i.p. injection [36].

**Fig. 5.**
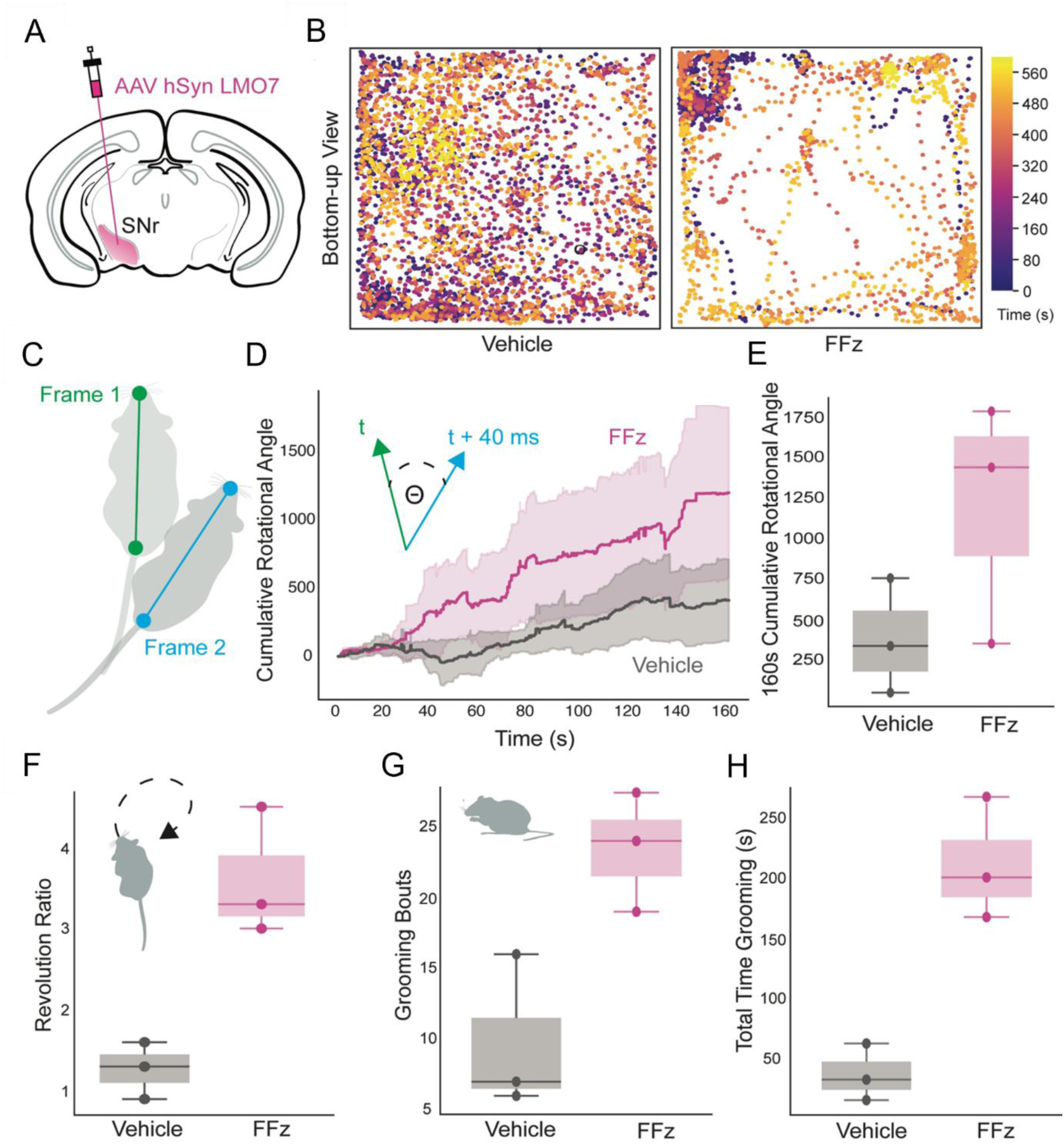
LMO7-mediated bioluminescence-driven excitation of SNr neurons elicits motor behavioral changes. (**A**) Schematic illustration of the expression strategy for unilateral photoactivation of substantia nigra pars reticulata (SNr) neurons. AAV-hSyn LMO7 was injected in SNr of wildtype mice. (**B**) Instantaneous time lapsed trajectory of the snout of a representative mouse in an open field treated with either vehicle (left) or FFz (right). (**C**) Schematic illustration of body parts (snout and base of the tail) extracted with DeepLabCut between consecutive video frames for instantaneous rotational analysis. (**D**) Summary data showing the cumulative rotation degrees in mice expressing LMO7 during a 160-second-long episode treated with vehicle or FFz. Inset, calculation of rotation degree (θ) for each frame (40 millisecond bins). n = 3 animals. Data are presented as mean values ± SEM. (**E**) Summary data showing the total degrees rotated in each experimental condition; p=0.04. (**F**) Turning preference displayed as the ratio of ipsilateral over contralateral revolutions during a 10 minute duration; p=0.01. (**G**) Total number of grooming bouts (left, p=0.06) and total time spent grooming (right, p=0.006) between vehicle and FFz treated mice.

When FFz was administered i.p., LMO7 expressing mice showed substantially greater rotational walking compared to vehicle injections (Fig. 5B-E). This effect was evident from the trajectories of the snout of the same mouse after vehicle versus FFz injection (Fig. 5B) and from plotting cumulative rotational angles of all three mice (Fig. 5C, D) 5–10 minutes after injection (160 seconds following the start of video recordings). Cumulative rotational angles were significantly different between vehicle and FFz treated mice (Fig. 5E, paired t-test, p = 0.04). As expected, a preference for ipsilateral rotation was detected compared to contralateral in FFz treated mice, as can be seen in the rotational ratios (Fig. 5F, paired t-test, p = 0.01). In addition to preference in body orientation, increased grooming was observed after unilateral LMO7 activation in SNr, presumably an effect of subthalamic nuclei projections [37]. FFz i.p. injections increased the duration of grooming significantly (Fig. 5G right panel, paired t-test, p = 0.006). We also note that a trend existed to an increased number of grooming bouts in FFz compared to vehicle (Fig. 5G left panel, paired t-test, p = 0.06).

## 4 Discussion

Here we describe the invention of a new excitatory LMO with improved efficiency of bioluminescent (chemogenetic) activation in response to small molecule luciferins (coelenterazines). We identified a novel luciferase variant obtained through molecular evolution of OLuc whose pairing with mNeonGreen resulted in bright FRET-based emission peaking at ∼518 nm. By tethering this emitter to VChR1 we generated a new LMO (LMO7) that outperforms our current standard (LMO3) in driving opsin activation. We show that, in presence of H-CTZ or FFz, LMO7 generates superior inward currents in HEK cells, promotes increased neural depolarization and spiking in brain slice and *in vivo* recordings, and can be used to control motor and grooming behavior in mice following viral injection.

We tested the ability of four different bioluminescent emitters (sbGluc, NanoLuc, NCS1 and NCS2) to drive bioluminescence-dependent activation of VChR1 in HEK cells. As compared to sbGluc and NanoLuc (both single luciferase emitters) we found that the two FRET probes (NCS1 and NCS2) had higher coupling efficiency and were able to generate larger inward currents, suggesting that, in addition to maximum brightness, close spectral pairing of emitter and opsin may be important at the comparably lower light intensity levels obtained with bioluminescence versus LED light sources. Whereas both FRET-probes featured emission peaks at ∼518 nm, the coupling efficiency of NCS2 to VChR1 proved to be significantly higher than for NCS1, suggesting that the orientation of NCS2 relative to the channel may also be favorable. As we observed substantially lower bioluminescent emission with LMO7 versus other LMOs, NCS2 may drive opsin activation more efficiently by facilitating higher levels of (non-radiant) FRET to the VChR1 chromophore.

Compared to previous versions of LMOs, LMO7 will allow more flexibility in experimental applications. Due to its higher coupling efficiency, LMO7 can be used with lower concentrations of luciferin, allowing i.p. application rather than intraventricular or intravenous delivery. Moreover, as the luciferase in LMO7 utilizes furimazine analogues, investigators can take advantage of the ongoing development of these variants that show improved brain access [22, 23, 36].

Several avenues exist for further optimization of this integrated control strategy. As brighter bioluminescent emitters and highly light sensitive opsins continue to be discovered and/or engineered [38, 39], and as new CTZ analogues with increased brain permeability, half-life and brightness are developed [23, 36], the ability to manipulate neural activity with biological light will continue to improve.

## Code, Data, and Materials Availability

Data supporting the article’s conclusions is available upon reasonable request to the corresponding author.

## Acknowledgments

We would like to thank all members of Bioluminescence Hub (http://www.bioluminescencehub.org/) laboratories for their feedback, discussions, and thoughtful comments throughout. This work was supported by the Swedish Research Council (AB: 2016-06760), National Institutes of Health grants (UH: R21MH101525, NCS: R01GM121944, and UH, NCS, CIM: U01NS099709) and National Science Foundation grants (UH, NCS: CBET-1464686 and CIM, DL, UH, NCS: DBI-1707352).

Conceptualization, AB, CIM, UH, NCS; Validation, AB, JM, ELC, GGL, CIM, UH, NCS; Formal Analysis, AB, JM, ELC; Investigation, AB, JM, ELC, ECI, GGL, MP, NF, TB; Data Curation, AB, JM, ELC; Writing Original Draft, AB, JM, ELC; Writing, Review & Editing, AB, NCS, CIM, UH; Visualization, AB, JM, ELC; Supervision, CIM, UH, NCS; Project Administration, UH; Funding Acquisition, AB, DL, CIM, UH, NCS.

