## Supplementary material for "A New Highly Efficient Molecule for Both Optogenetic and Chemogenetic Control Driven by FRET Amplification of BioLuminescence": Suppl. Figs 1-5

### **Supplementary Material** for Björefeldt *et al.* 2023

**Figures S1–S5.**

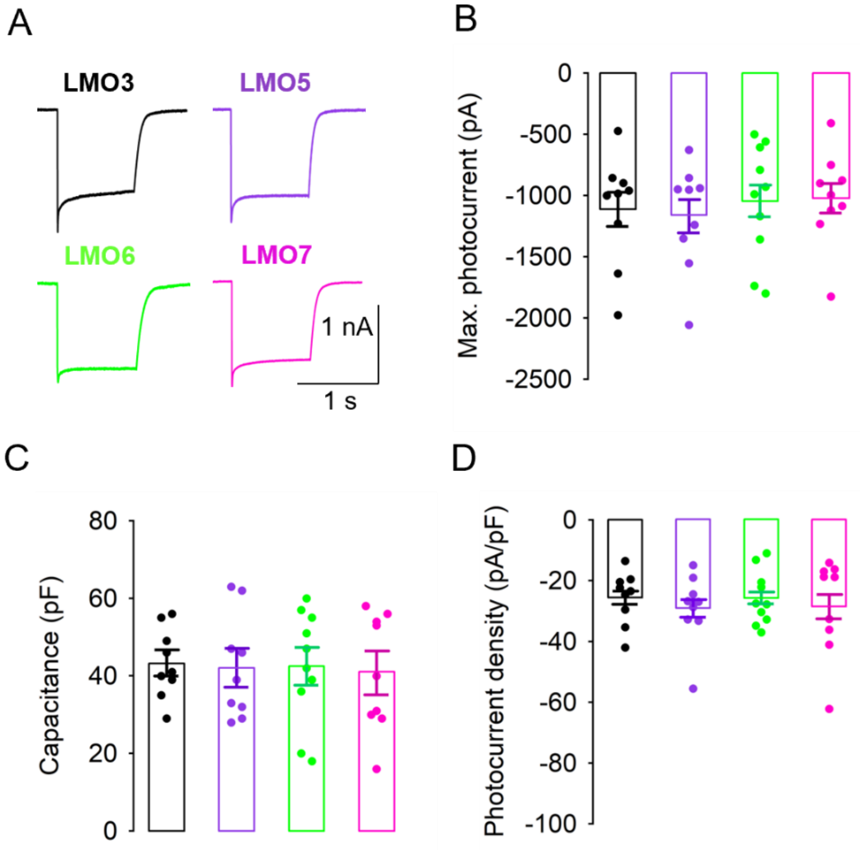


**Fig. S1** Average photocurrent amplitude and density across stable expression HEK lines. (**A**) Example traces of typical photocurrent amplitudes evoked by 1 second illumination at maximum irradiance. (**B**) Summary graph of average VChR1 photocurrent amplitudes evoked at maximum irradiance across HEK lines. (**C**) Summary graph showing capacitance values of all HEK cells and across stable lines. (**D**) Summary graph showing average photocurrent density across HEK lines.

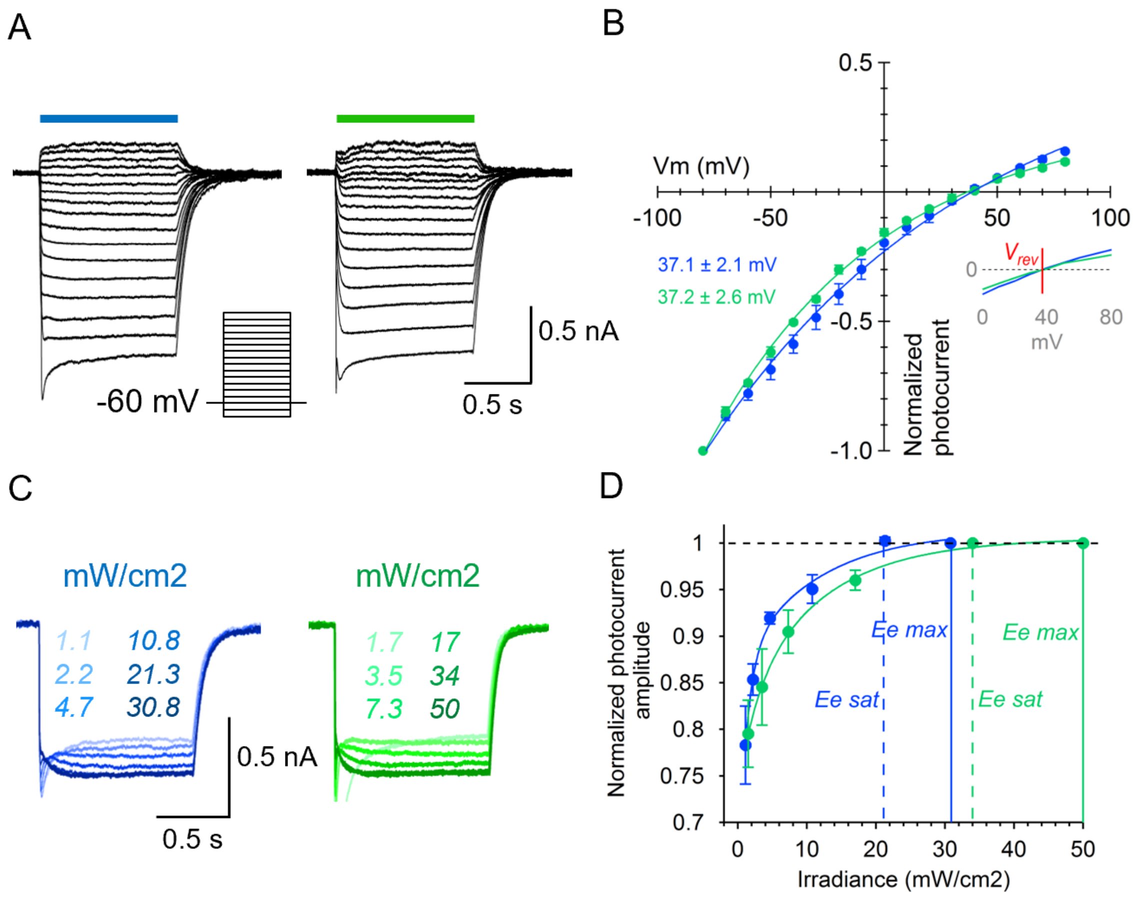


**Fig. S2** Characterization of photocurrent responses to blue and green light in HEK cells. (**A**) Example traces of photocurrent amplitudes evoked by 480 nm (left) and 540 nm (right) light in 1 second windows from -80 to +80 mV. (**B**) Summary graph of photocurrent-voltage relationship and reversal potential of VChR1 with blue and green light illumination (n = 4 cells from LMO3- and LMO7- expressing HEK lines). (**C**) Example traces of photocurrent amplitudes in response to increasing blue (left) and green (right) light intensity as indicated (**D**) Summary graph showing photocurrent amplitude relative to irradiance level with blue and green light (n = 3 LMO3 cells). Ee max; maximum irradiance level obtained, Ee sat; irradiance level sufficient to saturate photocurrent amplitude. Saturation occurs below maximum irradiance levels for both blue and green light.

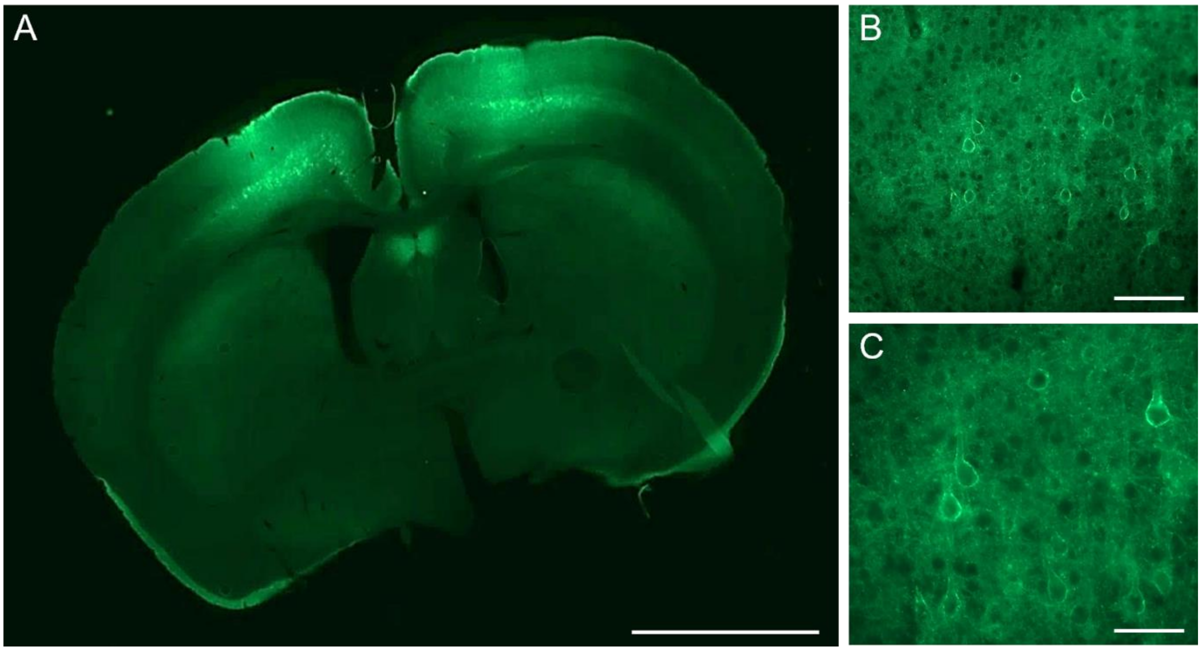


**Fig. S3** Expression of LMO7 in mouse neocortex after early postnatal bilateral AAV injection. (**A**) Representative image showing the distribution of LMO7 (mNeonGreen) fluorescence at 4 weeks post AAV9-hSyn-LMO7 injection into the fronto-paritetal cortex of P2 Swiss Webster mice. (**B**, **C**) magnified images from ‘A’ showing LMO7-expressing neurons at 20x (**B**) and 40x (**C**). Scale bars; A: 2 mm, B: 200 µm, C: 100 µm.

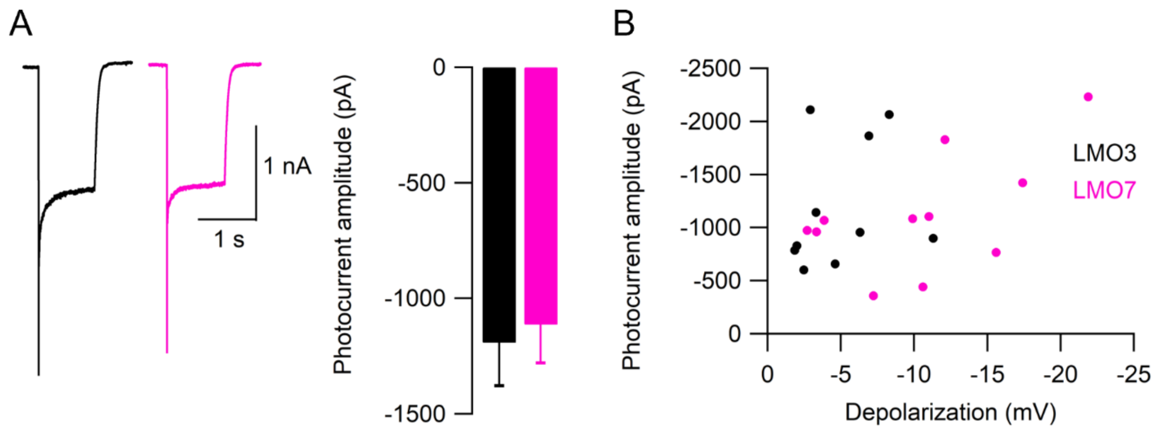


**Fig. S4** Average photocurrent amplitudes recorded from LMO3 and LMO7-expressing neurons. (**A**) representative photocurrent traces (left) and average photocurrent amplitudes (right) recorded from LMO3 (black) and LMO7 (magenta) populations (480 nm excitation light, 1s). (**B**) scatter plot showing photocurrent amplitude relative to CTZ-mediated depolarization for each recorded cells (LMO3; n = 10, LMO7; n = 11).

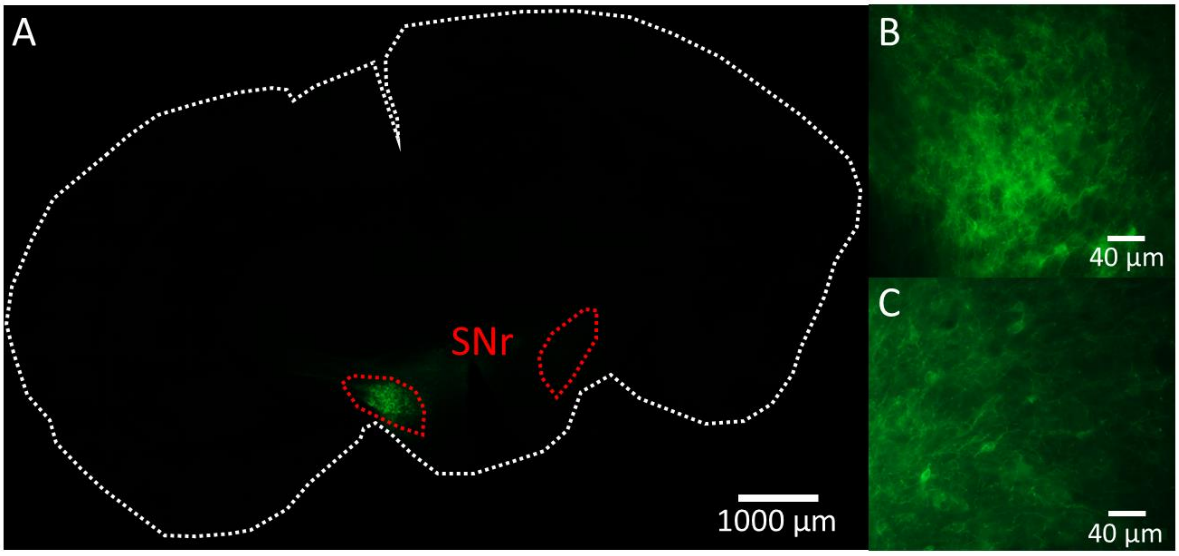


**Fig. S5** Expression of LMO7 in mouse substantia nigra pars reticulata (SNr) after unilateral AAV injection. (A) Representative image showing the localized expression of LMO7 (mNeonGreen) fluorescence at 4 weeks post AAV9-hSyn-LMO7 injection into the left SNr of C57BL/6 adult mice. White stippled line indicates section margins; red stippled lines mark area of SNr on both sites. (B, C) Magnified images from ‘A’ showing LMO7-expressing neurons at 20x from the center (B) and the periphery (C) of the labeled region.
